# A Genome-Wide Association Study of Non-Photochemical Quenching in response to local seasonal climates in *Arabidopsis thaliana*

**DOI:** 10.1101/539379

**Authors:** Tepsuda Rungrat, Andrew A. Almonte, Riyan Cheng, Peter J. Gollan, Tim Stuart, Eva-Mari Aro, Justin O. Borevitz, Barry Pogson, Pip B. Wilson

**Author notes:** These authors contributed equally to this work (Tepsuda Rungrat, Andrew A. Almonte).

## Abstract

Field-grown plants have variable exposure to sunlight as a result of shifting cloud-cover, seasonal changes, canopy shading, and other environmental factors. As a result, they need to have developed a method for dissipating excess energy obtained from periodic excessive sunlight exposure. Non-photochemical quenching (NPQ) dissipates excess energy as heat, however the physical and molecular genetic mechanics of NPQ variation are not understood. In this study, we investigated the genetic loci involved in NPQ by first growing different *Arabidopsis thaliana* accessions in local and seasonal climate conditions, then measured their NPQ kinetics through development by chlorophyll fluorescence. We used genome-wide association studies (GWAS) to identify 15 significant quantitative trait loci (QTL) for a range of photosynthetic traits, including a QTL co-located with known NPQ gene *PSBS* (AT1G44575). We found there were large alternative regulatory segments between the *PSBS* promoter regions of the functional haplotypes and a significant difference in PsbS protein concentration. These findings parallel studies in rice showing recurrent regulatory evolution of this gene. The variation in the *PSBS* promoter and the changes underlying other QTLs could give insight to allow manipulations of NPQ in crops to improve their photosynthetic efficiency and yield.

*B.P. & J.B. conceived the project; B.P., J.B., P.W. and T.R. designed the research plan and analysis; P.W. supervised the experiments; T.R. performed most of the experiments and analysis; P.G., T.S., A.A. & E.A. designed and undertook experimental design, experiments and analysis for Figure 4; R.C. did the GWAS analysis; P.W., T.R. & A.A. wrote the article with contributions of all the authors.*

## INTRODUCTION

Photosynthesis is the process of harnessing light energy to power CO_2_ fixation. Variability in light environments of a plant or leaf caused by clouds or canopy shading can often result in rapid switching between limited or excess light exposure in relation to the acclimated state. Excess light can result in the production of damaging reactive oxygen species, which can impair leaf photosynthesis even after return to low light. Plants have evolved a method to dissipate excess light energy from the major light-harvesting antennae as heat through a photoprotective mechanism known as non-photochemical quenching (NPQ; Reviewed in Ruban, 2016). This process can eliminate over 75% of absorbed light energy (Niyogi, Grossman, & Bjorkman, 1998) and modified regulation of NPQ has been shown to improve photosynthetic efficiency and increase biomass in field-grown tobacco (Kromdijk et al., 2016).

In vascular plants, the NPQ mechanism is generally classified into two main components: energy-dependent quenching (qE) and photoinhibition quenching (qI; Ruban, 2016). qE is recognized as the most significant component of NPQ and can be rapidly increased and relaxed within seconds to minutes. qE is triggered by a decrease in the thylakoid lumen pH (Briantais, Vernotte, Picaud, & Krause, 1979), which induces thermal dissipation through protonation of the PsbS protein, as well as the serial de-epoxidation of violaxanthin to antheraxanthin and then zeaxanthin through the xanthophyll cycle (Reviewed in Demmig-Adams & Adams, 1992; Jahns & Holzwarth, 2012). Conformational changes in the light-harvesting antennae activated during qE involve monomerization of PsbS (Correa-Galvis, Poschmann, Melzer, Stuhler, & Jahns, 2016; X. P. Li et al., 2004); however, the precise role of PsbS in the dissipation of excess energy is as yet unclear.

The worldwide Arabidopsis collection contains natural diversity in a range of traits that have resulted from adaptation to a wide range of climate types, and significant variation in NPQ was observed in a study of 62 diverse Arabidopsis accessions (Jung & Niyogi, 2009). The mapping of biparental populations from contrasting accessions has identified several novel loci involved in NPQ, including reinforcing the role of *PSBS* (Jung & Niyogi, 2009). However, to our knowledge, no studies have undertaken genome wide association studies (GWAS) on NPQ in Arabidopsis. GWAS surveys a much wider range of genetic diversity than bi-parental populations and also provides greater resolution for mapping quantitative trait loci (QTL) to assist in gene discovery. The utility of GWAS for NPQ has already been demonstrated through the characterization of 33 QTL in rice (Wang et al., 2017) and 15 QTL in soybean (Herritt, Dhanapal, & Fritschi, 2016).

The use of high-throughput phenotyping platforms that measure photoprotective traits using chlorophyll fluorescence have proven to be useful methods for monitoring real-time plant stress responses in model species such as Arabidopsis (Rousseau et al., 2013; Rungrat et al., 2016; van Rooijen, Aarts, & Harbinson, 2015). Phenotyping platforms such as PlantScreen can measure large numbers of plants simultaneously to reveal the photosynthetic performance of whole rosettes. Additionally, modified growth chambers can be used to precisely mimic external climates (Brown et al., 2014) without the noise of field conditions, facilitating the assessment of the impact of genetic variations on responses to even minor environmental variation. Combined with the extensive genetic resources available for Arabidopsis (1001 Genomes Consortium, 2016; Y. Li, Huang, Bergelson, Nordborg, & Borevitz, 2010; Zhang, Hause, & Borevitz, 2012), these tools enable the dissection of the genetic architecture underlying important photoprotective traits such as NPQ and their response to different environmental conditions.

In this study, we focus on identifying the effect of contrasting climates (in the form of differing light intensities and temperature profiles) on the kinetics of NPQ in several Arabidopsis accessions from the global diversity set. We then use GWAS to reveal the genetic basis underlying NPQ and its response to the environment. With these methods, we aim to better understand the genetic framework of this important physiological pathway in its response to excess light energy that occurs in natural environments.

## MATERIAL AND METHODS

### Plant growth

For GWAS, 284 genetically diverse Arabidopsis accessions were selected from the global HapMap set (Y. Li et al., 2010). Two photoprotective mutants (*npq1* and *npq4*; Y. Li et al., 2010; Niyogi et al., 1998) were included as controls for reduced NPQ. *npq1* is a loss of function mutant in violaxanthin de-epoxidase 1 (VxDE; AT1G08550) while *npq4* is a loss of function mutant in *PSBS* (AT1G44575). Most accessions had one replicate per environmental condition while 16 replicates of Col-0 were included in each condition to monitor the extent of spatial variation within the chamber and four replicates of *npq4* were included in the late autumn conditions. Seed germination was synchronised by stratification at 4°C in the dark in sterilised water for 4-5 days. Plants were grown in pots (4 cm × 4 cm × 7 cm) of pasteurised seed raising mix (Debco seed raising mix, Scotts Australia) without further fertilisation in specially modified climate chambers (Brown et al., 2014) housed in the plant growth facility of the Australian Plant Phenomics Facility at the ANU. These chambers have been fitted with 7-bands LED light panels and are programmed to alter light intensity, light spectrum, air temperature and relative humidity every 5 minutes. Climatic conditions were modelled using SolarCalc software (Spokas & Forcella, 2006). In this study, two experiments were run with diurnal and seasonal temperature fluctuations with two climates in each experiment set to simulate coastal (Wollongong: −34.425, 150.893) and inland (Goulburn: −34.426, 150.892) regions of South East Australia. The first experiment was conducted by simulating a typical late-autumn season starting from April 1st, 2014 and ending on June 5th, 2014. The second experiment was conducted to simulate an early autumn season, starting from March 15th, 2015 and finishing on May 7th, 2015. The maximum light intensity at noon was around 150 μmol photons m^−2^s^−1^ and 300 μmol photons m^−2^s^−1^ for Coastal and Inland, respectively, in both experiments. These are typical light intensities for growing *Arabidopsis thaliana* (Bölter, Seiler, & Soll, 2018). The temperature ranged between 14°C - 23°C and 10°C - 23°C for Coastal-Late Autumn and Inland-Late Autumn respectively, and these temperatures decreased to 10°C - 18°C and 5°C - 15°C by the end of the experiment due to seasonal change to winter. The temperature ranged between 10°C - 25°C and 5°C - 25°C for Coastal-Early Autumn and Inland-Late Autumn respectively, and these temperatures decreased to 10°C - 1°C and 5°C - 18°C by the end of the experiment due to seasonal change to winter.

### NPQ quantification by chlorophyll fluorescence measurement

Photosynthesis parameters were measured by pulse amplitude modulation chlorophyll fluorescence using the automated PlantScreen system (Photon Systems Instruments, Czech Republic) when plants were at 25 and 40 days of age and at 14 and 16 leaves stages. All measurements began at midday and finished by 5:00 pm. Chlorophyll fluorescence kinetics were monitored during illumination of actinic light (700 μmol m^−2^s^−1^) and saturation flashes (800ms, 2800 μmol m^−2^s^−1^) and analysed using FluorCam7 software. To focus on the major qE component, a custom-made chlorophyll fluorescence measurement protocol was used (P3; Rungrat et al., 2016). After 30 minutes dark adaptation, F_o_ was measured in the dark before F_m_ was measured with an initial saturating pulse in the dark, followed by a series of saturating pulses 60 seconds apart to monitor F_m_’ and F’ during the 8 minutes of actinic light illumination and following 3 minutes dark relaxation period. NPQ can be calculated from the ratio of change in F_m_ and F_m_’ during the illumination as shown in the equation:

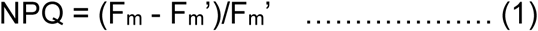

with F_m_’ and F_m_ being the maximal fluorescence of the light-adapted and dark-adapted leaf, respectively. In addition to NPQ, dark adapted F_v_/F_m_ (QY-max) was measured as an indicator of photoinhibition and light adapted F_v_’/F_m_’ was the measure of photosynthetic efficiency under the actinic light.

### Statistical analysis and QTL mapping

#### Experimental Design

Our goal is to identify quantitative trait loci as associated SNPs underlying photoprotection traits. In genome wide association studies (dominated by human genetics) the level of replication is at the level of the SNP, which ideally would be independent of other loci. Because of both local linkage disequilibrium and background population structure SNPs are not independent. This is particularly true with inbred accessions. Fortunately, a large collection of accessions is available which were selected to be roughly equidistantly related to each other (Platt et al, 2010). Thus, to maximise statistical power for identifying genetic variants underlying phenotypes, a panel of 284 natural accessions of Arabidopsis was selected from the HapMap collection for the Late Autumn experiment and an overlapping panel of 223 accessions was selected for the Early Autumn experiment. To maximize the number of accessions within an experiment, single replicates were grown in each climate condition. The Col-0 accession was replicated 16 times to allow calculation of biological variation of the traits.

For the phenotypic data, three phases of NPQ kinetics were examined. Induction was determined by NPQ formation when actinic light illumination was initiated, followed by a steady state phase and lastly, a relaxation phase in the dark (induction: 0 - 120 sec, steady state: 120 - 450 sec, and relaxation phases: 450 - 610 sec, Fig 2). The rate of induction was calculated as average slope of increase in NPQ during the induction (0 - 120 seconds); the maximum NPQ value was determined as the maximum value reached throughout the whole experiment and the rate of relaxation was calculated as the average slope during the dark relaxation (450 - 610 seconds). As relatively few, equally spaced, time points were sampled during NPQ, fitting nonlinear induction and relaxation kinetics is unlikely to change the results substantially. This is because the QTL effect is not the kinetics of particular accession, but the relative difference between summaries of the kinetic paths among SNP genotypes (Fig1).

**Figure 1:**
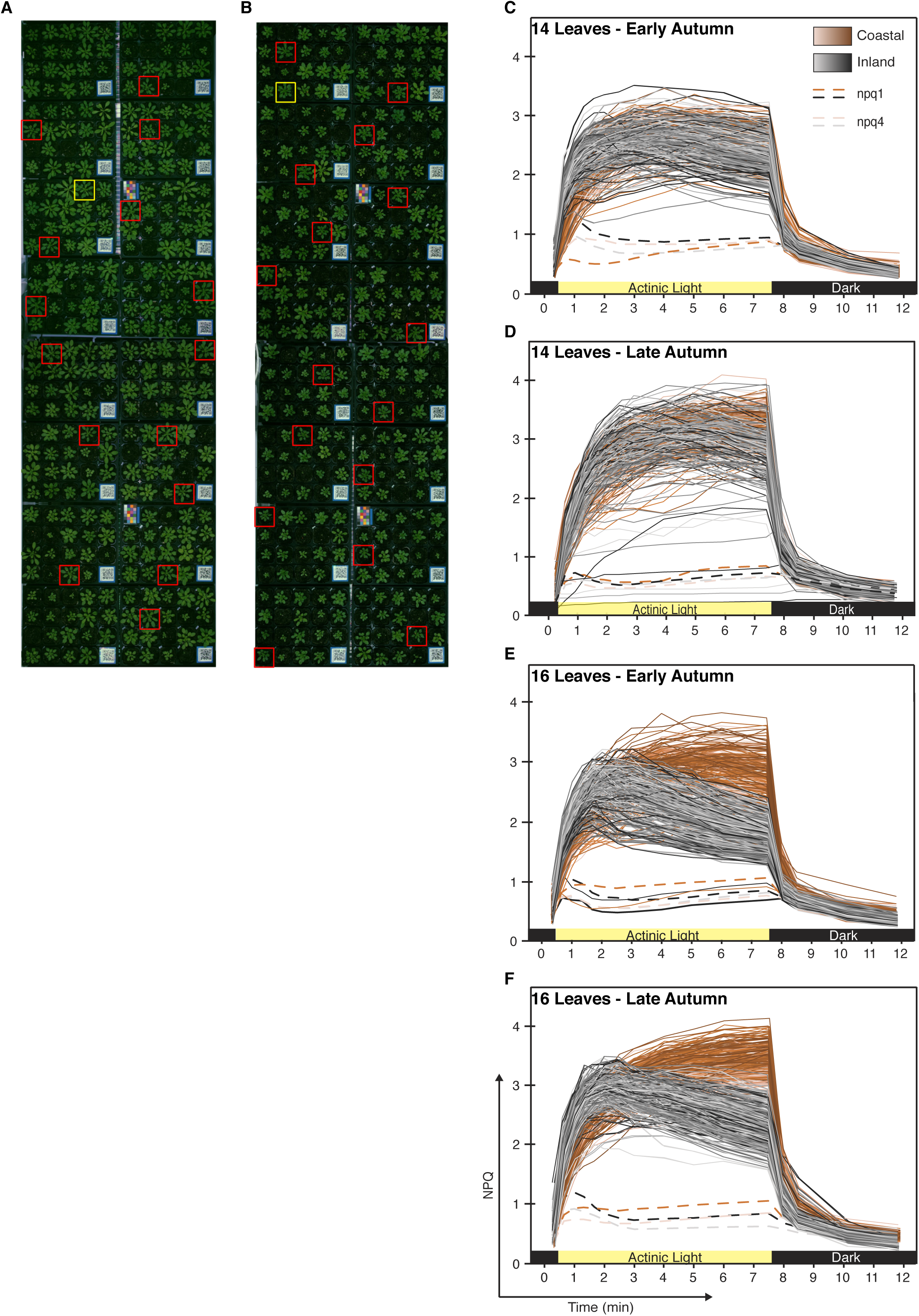
Natural variation of growth in response to different environmental conditions of 284 Arabidopsis accessions and 2 photoprotective mutants (*npq1*, *npq4*) within coastal (A) and inland (B) conditions. In this experiment, plants were measured for NPQ when Col-0 control plants (red boxes) reached the 16-leaf stage in both growth conditions to minimise developmental effects. (C – F) NPQ kinetic profiles of 14-leaf plants grown in early (C) and late (D) autumn conditions and 16-leaf plants grown in early (E) and late (F) autumn conditions.

**Figure 2:**
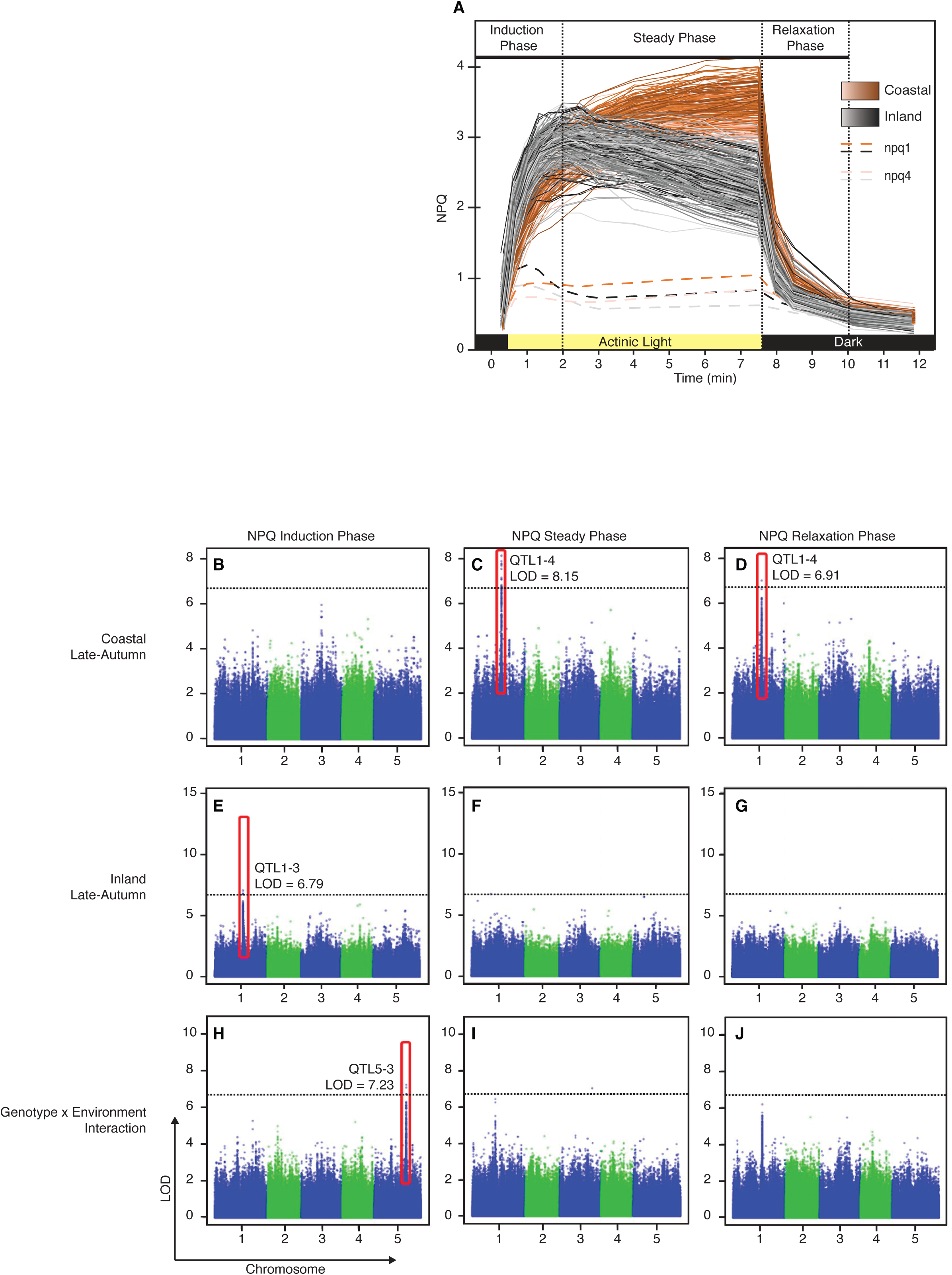
(A) NPQ kinetics profile of 16-leaf plants grown in late autumn conditions highlighting the key components of NPQ that were the focus of GWAS analysis. (B-J) Results of GWAS performed during the three stages of NPQ defined in (A) on plants grown in coastal late-autumn (B-D) and inland late-autumn (E-G) conditions, as well as Gene × Environment interactions (H-J) with the most significant SNPs highlighted. The dotted lines indicate the Bonferroni threshold of significance.

The individual phenotypes for each accession was used for the GWAS separately in both climate conditions. To determine the GxE interaction, all data was used in a single GWAS and each SNPs was tested for a main and climate specific effect and relatedness among accessions was accounted for as a random effect kinship matrix (Li et al, 2014). SNP data from the 6M SNPs data set (1001 Genomes Consortium, 2016) was filtered for a minor allele frequency <2.5% with a final set of approximately 1.7 million SNPs. GWAS were performed using the R package QTLRel (R. Cheng, Abney, Palmer, & Skol, 2011), based on model (2) below, which incorporated a relationship matrix to correct for confounding population structure.

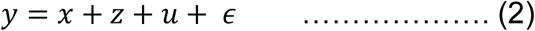

where *y* = (*y*_ij_) is a vector of the n phenotypic values with *y*_ij_ being the j-th accession in the i-th environment (i.e. one of the two climate conditions within each experiment), *x* = (*x*_ijk_)nx(K+1) represents the intercept and k covariates (if any) with effects β, *z* is a vector of the coded genotypes at a scanning locus with effect *γ*, *u* = (u_1_, u_2_, ……, u_n_)’ represents polygenic variation, and ∊ = (∊_1_, ∊_2_, ……, ∊_n_) the residual effect. It was assumed that 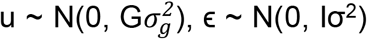 and *u* was independent of ∊. The genetic relationship matrix G was estimated by identify-by-state (IBS) from genotypic data with markers on the chromosome under scan being excluded to avoid proximal contamination (Riyan Cheng, Parker, Abney, & Palmer, 2013; Listgarten et al., 2012). A 0.05 genome-wide significance threshold was determined by the Bonferroni procedure, i.e., 2(1-0.05/m,1), where m is the number of SNPs. This threshold turned out to be very close to empirical thresholds estimated by the permutation test.

### Visualisation of alignments between KBS-Mac-74 and Col-0 accessions

Kablammo (Wintersinger & Wasmuth, 2015) was used to obtain a graphical understanding of the alignments between the *PSBS* genomic regions of the TAIR 10 Col-0 reference genome (Chr1: 16,866,832..16,873,428) and the recently sequenced KBS-Mac-74 genome (Michael et al., 2018). The sequence for the Col-0 *PSBS* genomic fragment was downloaded from the 1001 genomes project website and the KBS-Mac-74 genome was available on the European Nucleotide Archive (1001 Genomes Consortium, 2016; Michael et al., 2018). The KBS-Mac-74 genome was aligned with the Col-0 genomic fragment using Sequence Server v1.0.9 (Priyam et al., 2015) and the results were uploaded onto and visualised with Kablammo.

### Alignment of reads to Col-0 and KBS-Mac-74 haplotypes

Paired-end sequence reads for high and low NPQ accessions sequenced as part of the 1001 Genomes Project (1001 Genomes Consortium, 2016) were downloaded from NCBI SRA and trimmed for adapter content and low quality bases using Trimmomatic v0.36 (Bolger, Lohse, & Usadel, 2014) with the following parameters: LEADING:3 TRAILING:3 SLIDINGWINDOW:4:15 MINLEN:36. Retained paired-end reads were aligned to the Col-0 and KBS-Mac-74 genomes separately using Bowtie2 local alignment (1 mismatch per seed, maximum of 5 re-seeds; (Langmead & Salzberg, 2012). Alignment files were sorted and indexed using samtools (H. Li et al., 2009). Read coverage tracks were created for each accession using deepTools bamCoverage, with the coverage normalized to the number of reads per kilobase per million mapped reads (Ramirez et al., 2016). Coverage information around the *PSBS* genomic region was extracted from bedgraph files using bedtools v2.25 (Quinlan & Hall, 2010).

### Semi-quantification of PsbS protein in high and low NPQ lines

Five accessions with high NPQ and five accessions with low NPQ (Supplemental 6), as well as the *npq4* mutant and *PSBS* over-expression line, were grown for 6 weeks in 8 h day/16 h night cycle under 150 μmol photons m^−2^s^−1^ light with a temperature range of 8.6 °C – 18 °C. Mature leaves from 4 – 6 plants were pooled, frozen in liquid nitrogen, ground to a powder, and total proteins were extracted in 20 mM Tris-HCL solution (pH 7.8) containing 2 % SDS and protease inhibitor. After incubation at 37 °C and centrifugation, protein concentrations in the supernatants were measured using the Lowry assay. For each sample, 10 µg total protein were separated on SDS-PAGE gels containing 15% acrylamide, then transferred to PVDF membranes and blotted with a polyclonal antibody specific to the PsbS protein (a gift from R. Barbato). Six replicate Western blots containing all samples were developed and antibody signal intensities were quantified using an Odyssey CLx Imaging System (LI-COR). PsbS signals were normalised against a second non-specific protein band that was equally present in all samples (see Supplemental data). Statistical significance between the normalised signals was determined by a paired Student’s T test (N=30).

## RESULTS

### Natural variation of NPQ in Arabidopsis accessions in response to different climatic conditions

To determine the characteristics of NPQ in stressful environments, an Arabidopsis HapMap population of 284 accessions was grown in modified climate chambers (Brown et al., 2014) programmed to simulate contrasting environments: a “coastal” environment representing a temperate climate with light intensities similar to conditions conventionally used to grow Arabidopsis, and an “inland” environment representing a larger temperature range and higher light intensities (Supplemental 1). Two experiments were run with each condition starting at early- or late-autumn and transitioning into winter. There were clear differences in growth and development within the Arabidopsis HapMap population between the two conditions in both experiments (Fig 1A and B) as plants grown in inland conditions grew smaller than their coastal counterparts. Due to this variation in growth rate, environmental effects on NPQ phenotypes were measured when the Col-0 control plants reached a similar developmental stage (i.e. 14 or 16 leaves) rather than after a predefined period post-germination.

The NPQ kinetics of the plants showed variation dependent on their growth conditions and the developmental stage at which they were measured. When the samples were measured at the 14-leaf stage, plants grown in inland early-autumn conditions had a moderately faster NPQ induction relative to plants grown in coastal conditions by approximately 54%. However, the same plants grown in coastal and inland late-autumn conditions displayed largely similar NPQ kinetics (Fig 1C and D). All 14-leaf plants showed a rapid induction of NPQ that lasted approximately 1.5 – 2 minutes after exposure to actinic light (700 μmol m^−2^s^−1^), although plants grown in early-autumn conditions reached a maximum NPQ followed by a moderate decline throughout illumination (Fig 1C). In contrast, NPQ in plants grown in late-autumn conditions continued to increase during actinic light exposure (Fig 1D). Additionally, plants grown in late-autumn conditions reached a higher overall NPQ and returned more rapidly to basal levels during dark incubation following actinic light exposure.

NPQ kinetic profiles showed a dramatic difference between plants grown in coastal and inland conditions when measured at the 16-leaf growth stage (Fig 1E and F). Both groups had a rapid induction of NPQ within approximately 1.5 – 2 minutes of exposure to actinic light, though plants grown in coastal conditions were slightly slower to reach a steady phase. After the initial induction of NPQ upon actinic light exposure, plants grown in inland conditions reached their maximum average NPQ within 2 minutes, followed by a moderate and steady decrease in NPQ during illumination. Plants grown in coastal conditions continued to slowly increase during the same period before reaching their maximum NPQ after 7.5 minutes of illumination and directly before being transferred into darkness.

Of the plants grown to the 16-leaf stage, those grown in early-autumn conditions showed a larger variance in NPQ (Fig 1E – F). Inland plants measured in early-autumn achieved a maximum average NPQ of 2.47 (s.d. 0.38) after two minutes of exposure to actinic light, while coastal plants achieved a maximum average NPQ of 2.89 (s.d. 0.39) after five minutes of exposure. Plants measured in late-autumn conditions showed an overall higher NPQ, as inland plants achieved a maximum average NPQ of 2.62 (s.d. 0.88) after two minutes exposure to actinic light, while coastal plants achieved the highest NPQ average of 3.24 (s.d. 0.78) after 7.5 minutes of exposure to actinic light. When moved to darkness, the plants from inland conditions exhibited faster NPQ relaxation than plants grown in coastal conditions.

Photoprotective mutants *npq1*, lacking VxDE (Niyogi et al., 1998) and *npq4*, lacking PsbS (X. P. Li et al., 2000) were included as controls in all NPQ measurements. The mutants exhibited approximately four times lower steady-state NPQ than all wild-type accessions in both inland and coastal conditions and exhibited almost no NPQ relaxation (Fig 1C – F).

### Genome-wide association identifies 15 significant QTL for NPQ kinetics

In order to gain a better understanding of the QTL involved in NPQ, a GWAS was conducted using the R package QTLRel (R. Cheng et al., 2011) and the 6M SNP marker set (1001 Genomes Consortium, 2016). For mapping, a number of derived traits were calculated at both the 14- and 16-leaf stages, including the rate of NPQ induction, the slope of the steady phase, maximum NPQ value, and the rate of NPQ relaxation (Fig 2). For the late-autumn experiment, six QTL were identified across the three kinetic parameters of NPQ production, including QTL5-3 for the Genotype × Environment (GxE) interaction between the two conditions (Table 1, Fig 2H). For the early-autumn experiment, eight QTL were identified across the three kinetic parameters of NPQ production, including QTL2-3 and QTL4-1 for the GxE interaction. Interestingly, none of the NPQ QTL were identified in both experiments or in any two conditions. Mapping of the photosynthetic traits F_v_/F_m_‘ and QY-max under all conditions revealed five QTL, including QTL1-4 and QTL2-2 previously identified as NPQ QTL. The majority (12/15) of QTL were identified at the 16-leaf developmental stage, with only QTL2-2 being identified at both developmental stages but for different traits.

**Table 1.** Plants were grown in climatic growth chambers programmed to simulate the daily temperature ranges for late- and early-autumn conditions historically found in Australian coastal and inland environments. Temperature ranges gradually changed over time during the course of the experiments.

| Environment | Maximum light<br>intensity at noon<br>($\mu\text{mol m}^{-2} \text{s}^{-1}$) | Simulated seasons and dates | Temperature range | |
| --- | --- | --- | --- | --- |
|  |  |  | Beginning | End |
| Coastal | 150 | Early Autumn \| 15 Mar 2015 - 7 May 2015 | 14 - 24°C | 10 - 20°C |
|  |  | Late Autumn \| 1 Apr 2014 - 5 Jun 2014 | 13 - 23°C | 5 - 14°C |
| Inland | 300 | Early Autumn \| 15 Mar 2015 - 7 May 2015 | 11 - 24°C | 6 - 18°C |
|  |  | Late Autumn \| 1 Apr 2014 - 5 Jun 2014 | 10 - 23°C | 5 - 14°C |

**Table 2.** Fifteen QTL were identified for NPQ and other photosynthetic traits across four conditions and two developmental stages. The component of NPQ found to be associated with the specified QTL are listed under the conditions they were identified (colour-coded for convenience). G × E – genotype-by-environment interaction;^*e*^ analysis results at 14-leaf stage;^*f*^ analysis results at 16-leaf stage

| QTL | Chr | Position (bp) | LOD score | Col-0 allele Frequency | % variation explained | Trait appearance |  |  |  |  |  |
| --- | --- | --- | --- | --- | --- | --- | --- | --- | --- | --- | --- |
|  |  |  |  |  |  | Late Autumn |  |  | Early Autumn |  |  |
|  |  |  |  |  |  | Coastal | Inland | G x E | Coastal | Inland | G x E |
| QTL1-1 | 1 | 349 233 | 8.66 | 0.24 | 3.4 | NPQ Steady<sup>e</sup> |  |  |  |  |  |
| QTL1-2 | 1 | 9 729 376 | 7.04 | 0.36 | 9.2 | | | | $F_v'/F_m'^f$ | | |
| QTL1-3 | 1 | 16 351 441 | 7.04 | 0.52 | 5.6 |  | NPQ Slope Induction<sup>f</sup> |  |  |  |  |
| QTL1-4 | 1 | 16 869 695 | 8.15 | 0.79 | 9.5 | $F_v'/F_m'^f$ | QY-max<sup>f</sup> | | | | |
|  |  |  |  |  |  | NPQ Steady<sup>f</sup> |  |  |  |  |  |
|  |  |  |  |  |  | NPQ Max<sup>f</sup> |  |  |  |  |  |
|  |  |  |  |  |  | NPQ Slope Relaxation<sup>f</sup> |  |  |  |  |  |
| QTL1-5 | 1 | 20 680 932 | 7.93 | 0.05 | 13.7 |  |  |  |  | NPQ Steady<sup>f</sup> |  |
| QTL2-1 | 2 | 10 934 450 | 8.07 | 0.07 | 10.1 | | $F_v'/F_m'^f$ | | | | |
| QTL2-2 | 2 | 10 984 417 | 7.95 | 0.06 | 12.2 |  | NPQ Steady<sup>e</sup> |  |  |  |  |
|  |  |  |  |  |  |  | QY-max<sup>f</sup> |  |  |  |  |
| QTL2-3 | 2 | 11 680 235 | 7.00 | 0.21 | 6.7 |  |  |  |  |  | NPQ Slope Induction<sup>f</sup> |
| QTL4-1 | 4 | 1 509 645 | 7.95 | 0.19 | 0.9 | | | | | $F_v'/F_m'^f$ | $F_v'/F_m'^f$ |
| QTL4-2 | 4 | 5 270 810 | 7.14 | 0.22 | 7.5 |  |  |  |  | NPQ Slope Induction<sup>f</sup> |  |
| QTL4-3 | 4 | 13 911 309 | 9.13 | 0.06 | 0.1 |  |  |  | NPQ Steady<sup>e</sup> |  |  |
| QTL5-1 | 5 | 15 037 793 | 6.93 | 0.09 | 1.4 | NPQ Relaxation<sup>f</sup> |  |  |  |  |  |
| QTL5-2 | 5 | 15 930 795 | 7.27 | 0.22 | 11.7 |  |  |  |  | NPQ Relaxation<sup>f</sup> |  |
| QTL5-3 | 5 | 18 511 604 | 7.23 | 0.26 | 2.3 |  |  | NPQ Slope Induction<sup>f</sup> |  |  |  |
| QTL5-4 | 5 | 25 477 423 | 7.71 | 0.11 | 14.0 |  |  |  | NPQ Steady<sup>f</sup> |  |  |

These QTL were investigated in the TAIR database (Table 1; Huala et al., 2001), but a majority were not associated with an obvious candidate gene. QTL1-4 was a strong QTL identified in the coastal late-autumn condition for the slope of the steady phase and rate of NPQ relaxation (Fig 2C – D), as well as F_v_/F_m_ and maximum NPQ value (Table 1). It was also associated with QY-max in the inland late-autumn condition where it was slightly below the significance threshold. The SNP with the highest logarithm of odds (LOD) score for QTL1-4 was located in the promoter of the candidate gene *photosystem II subunit S* (*PSBS*; AT1G44575; Fig 3).

**Figure 3:**
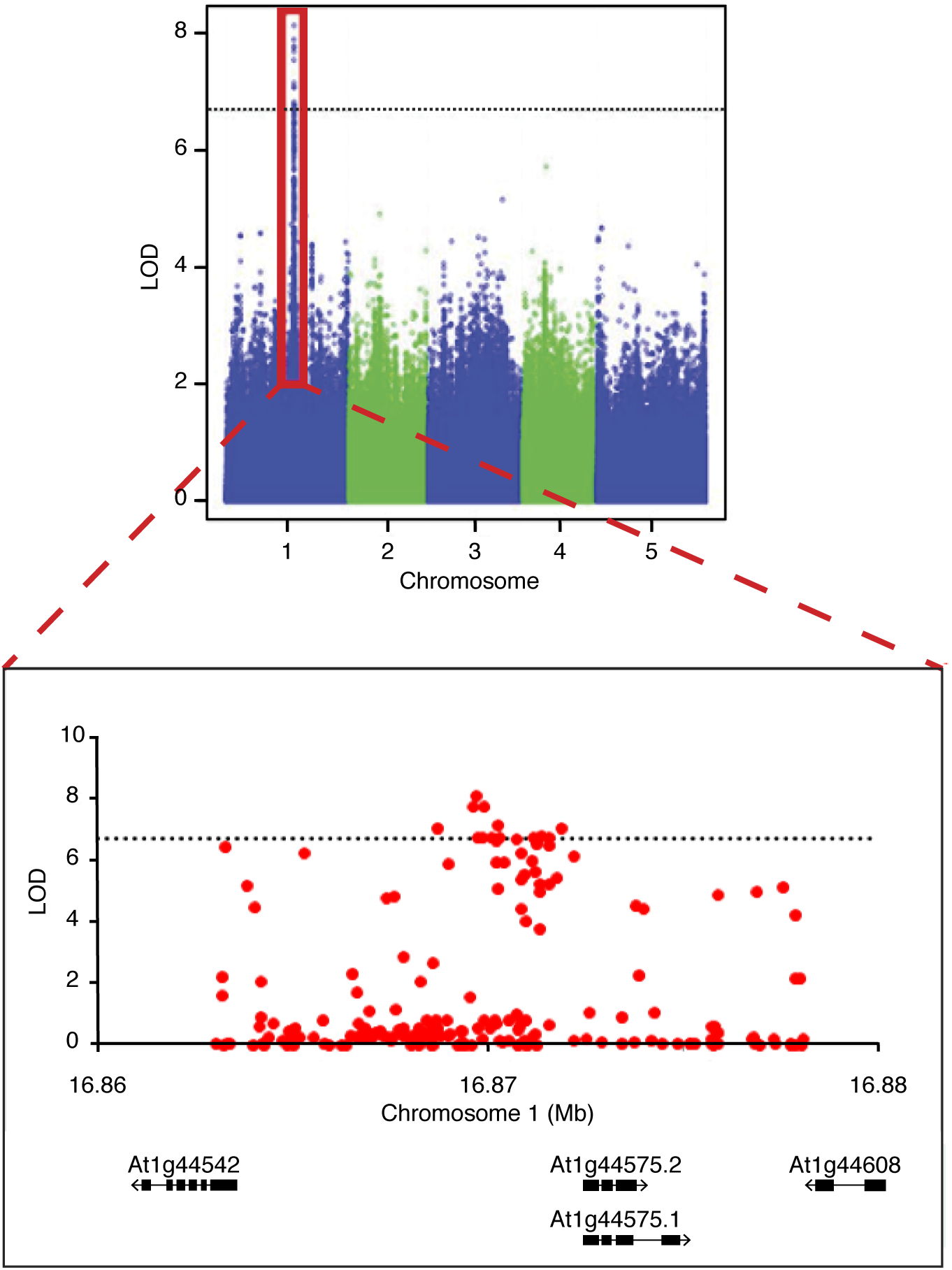
Association of SNPs in the region of QTL1-4, a QTL found to be significant during both the NPQ steady and relaxation phases for plants grown in coastal late-autumn conditions. The more significant SNPs are concentrated within the promoter region of the *PSBS* gene (AT1G44575). The black dotted lines indicate the Bonferroni threshold of significance.

**Figure 4:**
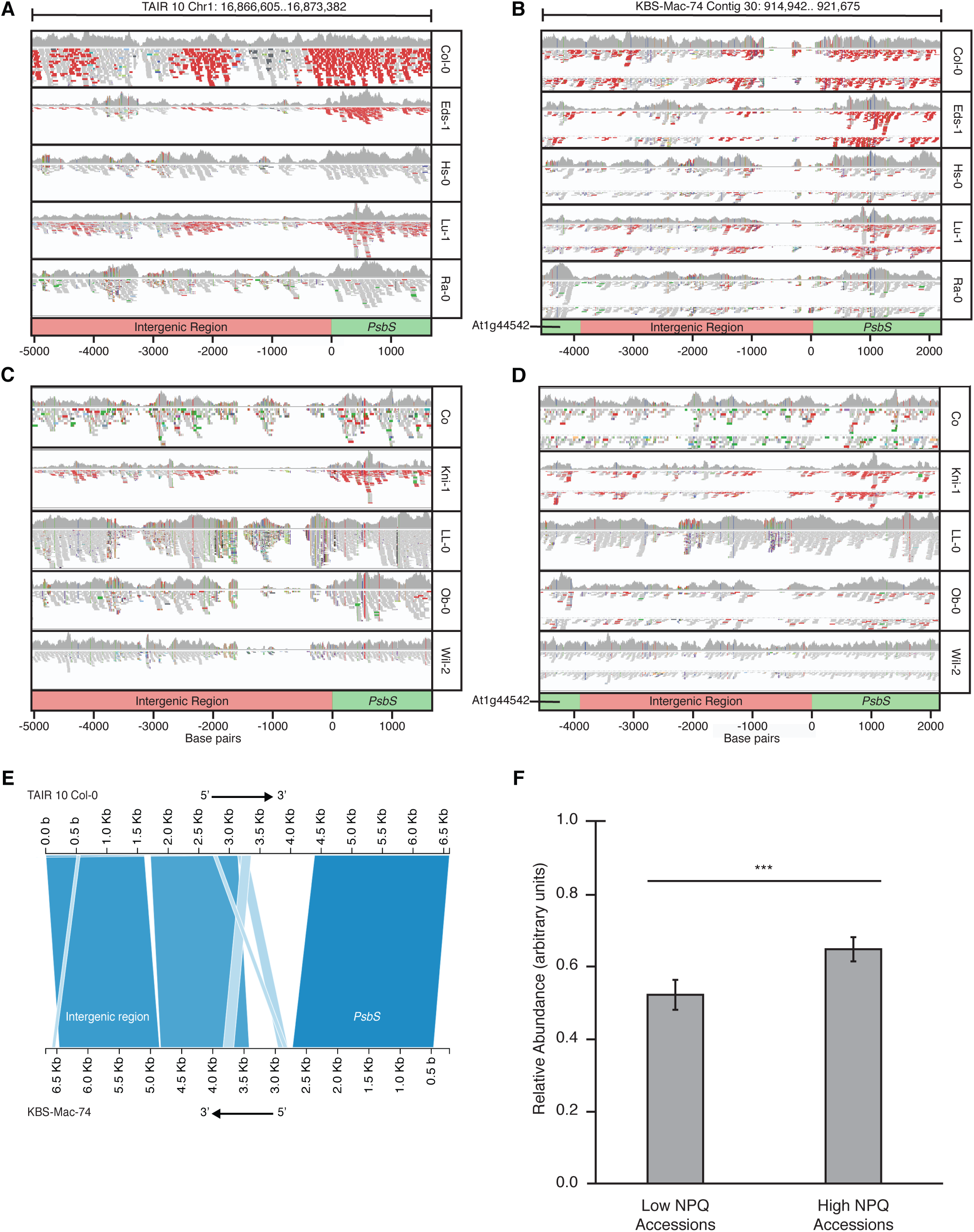
(A-D) Coverage tracks of the *PSBS* genomic regions from five low NPQ (A and B) and five high NPQ (C and D) Arabidopsis accessions aligned with the TAIR 10 Col-0 reference genome (low NPQ accession; A and C) and KBS-Mac-74 genome (high NPQ accession; B and D). Values along x-axes indicate the base pair distance relative to the *PSBS* transcription start site. Genes along the track are coloured green and the intergenic region is coloured pink. (E) Graphical view of the alignment of the TAIR 10 Col-0 and the KBS-Mac-74 *PSBS* genomic regions. Axis values refer to base pair positions within the respective tracks. (F) Comparison of the average relative PsbS protein abundance between low and high NPQ accessions. Error bars represent standard deviations. N=30; *** P < 0.001 with paired Student’s T test.

### Sequence variation in the PsbS promoter indicates differential induction of *PSBS* that may be important in NPQ regulation

To explore a possible relationship between natural variation in the *PSBS* genomic region and the diversity in NPQ phenotypes, the SNP-corrected sequences of the ten highest and ten lowest NPQ accessions determined in this experiment were acquired from the 1001 genomes project (Supplemental 1B; Weigel & Mott, 2009). Four SNPs were identified in the PsbS coding region in a comparison between the two haplotype groups (low NPQ and high NPQ), although each of these encoded synonymous amino acid residues (Supplemental 1A). Nevertheless, as these are SNP-corrected sequences, there may be major insertions or deletions that may not have been identified.

To compare the sequences of the high and low NPQ accessions, the raw reads of the *PSBS* genomic region from the 1001 genomes project were aligned to both the Col-0 reference genome (TAIR 10) and the recently sequenced KBS-Mac-74 genome (Michael et al., 2018). KBS-Mac-74 was chosen because it was the only accession available with long-read genomic sequence data and has not been SNP-corrected. Seven of the ten high NPQ accessions were available in the 1001 genomes project and all seven showed the same pattern of missing sections in the alignments to the promoter of *PSBS* to the Col-0 reference but aligned well to the KBS-Mac-74 genome. Similarly, of the eight low NPQ accessions available, all eight aligned well to the Col-0 reference but had missing sections in the alignment to the *PSBS* promoter from the KBS-Mac-74 genome (Fig 4A – D). When the Col-0 and KBS-Mac-74 genomes were aligned at this region there were two sections in the promoter of *PSBS* that did not align. One section was 1,251 bp in Col-0 corresponding with 691 bp in KBS-Mac-74, and the other was 100 bp in Col-0 corresponding with 3,000 bp in KBS-Mac-74 (Fig 4E).

To determine if there was a difference in PsbS protein abundance between high and low NPQ accessions, PsbS protein was quantified by Western blotting. Five plants representative of each NPQ haplotype were grown for six weeks under coastal late-autumn conditions. A small but highly significant difference in PsbS abundance between the two haplotype groups was identified (Fig 4F), with the high NPQ haplotype plants producing ~30% higher amounts of PsbS on average.

## DISCUSSION

In this study, climate chambers were used to investigate the effect of local seasonal growing conditions on the non-photochemical quenching (NPQ) pathway. A genome-wide association study (GWAS) was used to investigate the genetic architecture of NPQ and determine what quantitative trait loci (QTL) are involved in this physiological process, as it is important for high light tolerance.

It was clear that different climate conditions affected growth, as plants grown in coastal conditions grew larger than their inland cohort. The inland environment was programmed to provide higher light intensities and lower temperatures than the coastal environment, so NPQ would be more active in plants grown in inland conditions. Leaf expansion is inversely correlated with light intensity (Potter, Rood, & Zanewich, 1999) and content of the plant hormone gibberellic acid, which may underlie why plants grown in high-light environments are physically smaller (Reviewed in Hedden & Sponsel, 2015; Stowe & Yamaki, 1957).

The NPQ kinetic profiles of the plants in this study were found to be influenced by development and climate conditions. When grown to the 14-leaf stage, there was little difference in the NPQ kinetic profiles between plants grown in either inland or coastal climates. However, their NPQ profiles changed dramatically when grown to the 16-leaf stage. Despite these changes, there were some commonalities. For example, plants grown in inland conditions had consistently faster induction of NPQ when exposed to actinic light, which suggests the relatively more stressful growing environment has primed these plants to more rapidly respond to stressful conditions. Furthermore, all plants in all conditions rapidly resolved their NPQ when moved to darkness, though inland-grown 16-leaf plants did so more quickly. The steady decrease in NPQ observed in the inland-grown 16-leaf plants during their exposure to actinic light also suggests they are better acclimated to utilise abiotic stress response pathways. Differences in glutathione content between inland- and coastal-grown plants may be a potential mechanism for this observation as glutathione is a potent antioxidant that fluctuates with light intensity (Alsharafa et al., 2014). However, more research must be done to confirm this. The differences in NPQ kinetic profiles between plants grown under field-simulating high- and low-light environments demonstrates how dynamic environments can condition plants to develop appropriate physiological adaptations. Genetic variation in this response may underlie such adaptations to the particular light environments where the plants were collected.

To determine the genetic architecture of NPQ, a GWAS was performed and a total of 15 QTL were found to be involved in different components of NPQ (Table 1). They were detected across different developmental stages and different seasonal climatic conditions. The majority were found in the 16-leaf stage, further implying the relevance of plant development in local seasonal climates on NPQ efficiency. Arabidopsis has a wide geographic distribution and each accession would experience different conditions in their native environments. This variation was revealed through QTL that varied between the two simulated environments. All 15 QTL identified were associated with multiple candidate genes (Supplemental 7). For example, QTL1-5 may be associated with *NDF5*, a gene involved in regulation of gene expression and electron transport in photosystem I (Ishida et al., 2009). QTL5-2 may be associated with *PTST*, which encodes a chloroplast-localised protein that is involved in protein translocation (Lohmeier-Vogel et al., 2008).

QTL1-4 is the most prominent QTL identified in this study and is associated with *PSBS*, a gene known to be involved with NPQ (X. P. Li et al., 2000). The exact role of the PsbS protein in the NPQ pathway is currently unclear. It has been found to bind to chlorophylls and xanthophylls, making it the possible site of xanthophyll-dependent NPQ (X. P. Li et al., 2000). PsbS has also been found to induce conformational changes in the antennae of photosystem II (Horton, Wentworth, & Ruban, 2005). Regardless, PsbS is known to be a requirement because *npq4* is a PsbS loss-of-function mutant and consequently shows little to no NPQ activity (X. P. Li et al., 2000).

This study also found evidence for a correlation between NPQ competency and the *PSBS* genomic architecture. Gene alignments were performed for the highest and lowest NPQ accessions, with TAIR 10 Col-0 and KBS-Mac-74 (Michael et al., 2018) being the models for low and high NPQ accessions, respectively (Fig 4). These alignments show little variation within the protein-coding regions of *PSBS*, which ultimately have no impact on protein function since they do not alter the amino acid sequence. Alignments of the *PSBS* genomic regions from high and low NPQ accessions show structural variation within the promoter elements of the *PSBS* gene. The non-homologous elements may be due to a deletion, transposition, and/or a gene conversion event. There did not appear to be any obvious regulatory elements within those promoter regions, though more experimentation and functional tests would be required to determine their precise effect on PsbS expression and NPQ. Regardless, there does appear to be a significant influence on PsbS protein expression as high NPQ accessions have a relatively higher level of leaf PsbS protein content when grown in coastal late-autumn conditions (Fig 4F).

Besides genotypic and phenotypic variations in NPQ, there are also several instances of genotype-by-environment (GxE) interaction, mostly in NPQ induction. As previously mentioned, this may be due to plants grown in high-light environments being better conditioned to respond to light stress. QTL2-3 and QTL5-3 are both identified to have GxE variation in NPQ induction, but neither are associated with obvious candidate genes that may be directly or indirectly involved in NPQ. Nevertheless, a previous GWAS investigating the effects of climate on flowering time identified genes that would not have obviously been attributed to that pathway (Tabas–Madrid et al., 2018). A similar situation may be apparent in this study, but more research is required.

NPQ is an extremely important physiological pathway that allows plants to adapt to fluctuating lighting conditions and is therefore strongly influenced by the local environment. Examples of adaptations that result from such environmental pressures include the apparent increased reliance on the mechanism of the PsbS protein. This study has also shown there are several genetic components to NPQ identified by GWAS across simulated environments. Furthermore, there is evidence for a significant genomic event that directly impacts PsbS expression and subsequently impacts NPQ competence. In the face of current global climate challenges, better understanding of the NPQ pathway could have important implications for agriculture (Kromdijk et al., 2016) and may help better implement genomics-based strategies for food security and conservation efforts.

## ACKNOWLEDGEMENTS

This work was supported by grants from the ARC centre of excellence in Plant Energy Biology and the Australian National University for TR, AA, RC, PW, JO and BP and also for providing publication cost. The Australian Plant Phenomics Facility is supported under the National Collaborative Research Infrastructure Strategy of the Australian Government. TS is supported by the Jean Rogerson Postgraduate Scholarship. PG and E-MA acknowledge support from Academy of Finland projects 26080341, 307335 and 303757.

## REFERENCES

Alsharafa, K., Vogel, M. O., Oelze, M.-L., Moore, M., Stingl, N., König, K., … Dietz, K.-J. (2014). Kinetics of retrograde signalling initiation in the high light response of Arabidopsis thaliana. Phil. Trans. R. Soc. B, 369(1640), 20130424.

Bolger, A. M., Lohse, M., & Usadel, B. (2014). Trimmomatic: a flexible trimmer for Illumina sequence data. Bioinformatics, 30(15), 2114–2120. doi:10.1093/bioinformatics/btu170

Bölter, B., Seiler, F., Soll, J. (2018). Analysis of *Arabidopsis thaliana* Growth Behavior in Different Light Qualities. J. Vis. Exp. (132), e57152, doi:10.3791/57152.

Briantais, J. M., Vernotte, C., Picaud, M., & Krause, G. H. (1979). Quantitative Study of the Slow Decline of Chlorophyll Alpha-Fluorescence in Isolated-Chloroplasts. Biochimica Et Biophysica Acta, 548(1), 128–138. doi:Doi 10.1016/0005-2728(79)90193-2

Brown, T. B., Cheng, R. Y., Sirault, X. R. R., Rungrat, T., Murray, K. D., Trtilek, M., … Borevitz, J. O. (2014). TraitCapture: genomic and environment modelling of plant phenomic data. Current Opinion in Plant Biology, 18, 73–79. doi:10.1016/j.pbi.2014.02.002

Cheng, R., Abney, M., Palmer, A. A., & Skol, A. D. (2011). QTLRel: an R Package for Genome-wide Association Studies in which Relatedness is a Concern. Bmc Genetics, 12. doi:10.1186/1471-2156-12-66

Cheng, R., Parker, C. C., Abney, M., & Palmer, A. A. (2013). Practical considerations regarding the use of genotype and pedigree data to model relatedness in the context of genome-wide association studies. G3: Genes, Genomes, Genetics, g3. 113.007948. 1001

Genomes Consortiom (2016). 1,135 Genomes Reveal the Global Pattern of Polymorphism in Arabidopsis thaliana. Cell, 166(2), 481–491. doi:10.1016/j.cell.2016.05.063

Correa-Galvis, V., Poschmann, G., Melzer, M., Stuhler, K., & Jahns, P. (2016). PsbS interactions involved in the activation of energy dissipation in Arabidopsis. Nature Plants, 2(2). doi:10.1038/Nplants.2015.225

Demmig-Adams, B., & Adams, W. W. (1992). Photoprotection and Other Responses of Plants to High Light Stress. Annual Review of Plant Physiology and Plant Molecular Biology, 43, 599–626. doi:DOI 10.1146/annurev.pp.43.060192.003123

Hedden, P., & Sponsel, V. (2015). A century of gibberellin research. Journal of plant growth regulation, 34(4), 740–760.

Herritt, M., Dhanapal, A. P., & Fritschi, F. B. (2016). Identification of Genomic Loci Associated with the Photochemical Reflectance Index by Genome-Wide Association Study in Soybean. Plant Genome, 9(2). doi:10.3835/plantgenome2015.08.0072

Horton, P., Wentworth, M., & Ruban, A. (2005). Control of the light harvesting function of chloroplast membranes: The LHCIId-aggregation model for non-photochemical quenching. Febs Letters, 579(20), 4201–4206.

Huala, E., Dickerman, A. W., Garcia-Hernandez, M., Weems, D., Reiser, L., LaFond, F., … Huang, W. (2001). The Arabidopsis Information Resource (TAIR): a comprehensive database and web-based information retrieval, analysis, and visualization system for a model plant. Nucleic Acids Research, 29(1), 102–105.

Ishida, S., Takabayashi, A., Ishikawa, N., Hano, Y., Endo, T., & Sato, F. (2009). A novel nuclear-encoded protein, NDH-dependent cyclic electron flow 5, is essential for the accumulation of chloroplast NAD (P) H dehydrogenase complexes. Plant and cell physiology, 50(2), 383–393.

Jahns, P., & Holzwarth, A. R. (2012). The role of the xanthophyll cycle and of lutein in photoprotection of photosystem II. Biochimica et Biophysica Acta (BBA)-Bioenergetics, 1817(1), 182–193.

Jung, H. S., & Niyogi, K. K. (2009). Quantitative Genetic Analysis of Thermal Dissipation in Arabidopsis. Plant Physiology, 150(2), 977–986. doi:10.1104/pp.109.137828

Kromdijk, J., Glowacka, K., Leonelli, L., Gabilly, S. T., Iwai, M., Niyogi, K. K., & Long, S. P. (2016). Improving photosynthesis and crop productivity by accelerating recovery from photoprotection. Science, 354(6314), 857–861. doi:10.1126/science.aai8878

Langmead, B., & Salzberg, S. L. (2012). Fast gapped-read alignment with Bowtie 2. Nature Methods, 9(4), 357–U354. doi:10.1038/Nmeth.1923

Li, Y., Cheng, R., Spokas, K.A., Palmer, A.A., & Borevitz, J.O. (2014). Genetic Variation for Life History Sensitivity to Seasonal Warming in Arabidopsis thaliana. Genetics, 196(2): 569–577

Li, H., Handsaker, B., Wysoker, A., Fennell, T., Ruan, J., Homer, N., … Proc, G. P. D. (2009). The Sequence Alignment/Map format and SAMtools. Bioinformatics, 25(16), 2078–2079. doi:10.1093/bioinformatics/btp352

Li, X. P., Bjorkman, O., Shih, C., Grossman, A. R., Rosenquist, M., Jansson, S., & Niyogi, K. K. (2000). A pigment-binding protein essential for regulation of photosynthetic light harvesting. Nature, 403(6768), 391–395. doi:Doi 10.1038/35000131

Li, X. P., Gilmore, A. M., Caffarri, S., Bassi, R., Golan, T., Kramer, D., & Niyogi, K. K. (2004). Regulation of photosynthetic light harvesting involves intrathylakoid lumen pH sensing by the PsbS protein. Journal of Biological Chemistry, 279(22), 22866–22874. doi:10.1074/jbc.M402461200

Li, Y., Huang, Y., Bergelson, J., Nordborg, M., & Borevitz, J. O. (2010). Association mapping of local climate-sensitive quantitative trait loci in Arabidopsis thaliana. Proceedings of the National Academy of Sciences of the United States of America, 107(49), 21199–21204. doi:10.1073/pnas.1007431107

Listgarten, J., Lippert, C., Kadie, C. M., Davidson, R. I., Eskin, E., & Heckerman, D. (2012). Improved linear mixed models for genome-wide association studies. Nature Methods, 9(6), 525.

Lohmeier-Vogel, E. M., Kerk, D., Nimick, M., Wrobel, S., Vickerman, L., Muench, D. G., & Moorhead, G. B. (2008). Arabidopsis At5g39790 encodes a chloroplast-localized, carbohydrate-binding, coiled-coil domain-containing putative scaffold protein. BMC Plant Biology, 8(1), 120.

Michael, T. P., Jupe, F., Bemm, F., Motley, S. T., Sandoval, J. P., Lanz, C., … Ecker, J. R. (2018). High contiguity Arabidopsis thaliana genome assembly with a single nanopore flow cell. Nature communications, 9(1), 541.

Niyogi, K. K., Grossman, A. R., & Bjorkman, O. (1998). Arabidopsis mutants define a central role for the xanthophyll cycle in the regulation of photosynthetic energy conversion. Plant Cell, 10(7), 1121–1134. doi:DOI 10.1105/tpc.10.7.1121

Potter, T. I., Rood, S. B., & Zanewich, K. P. (1999). Light intensity, gibberellin content and the resolution of shoot growth in Brassica. Planta, 207(4), 505–511.

Priyam, A., Woodcroft, B. J., Rai, V., Munagala, A., Moghul, I., Ter, F., … Rumpf, W. (2015). Sequenceserver: a modern graphical user interface for custom BLAST databases. bioRxiv, 033142.

Quinlan, A. R., & Hall, I. M. (2010). BEDTools: a flexible suite of utilities for comparing genomic features. Bioinformatics, 26(6), 841–842. doi:10.1093/bioinformatics/btq033

Ramirez, F., Ryan, D. P., Gruning, B., Bhardwaj, V., Kilpert, F., Richter, A. S., … Manke, T. (2016). deepTools2: a next generation web server for deep-sequencing data analysis. Nucleic Acids Research, 44(W1), W160–W165. doi:10.1093/nar/gkw257

Rousseau, C., Belin, E., Bove, E., Rousseau, D., Fabre, F., Berruyer, R., … Boureau, T. (2013). High throughput quantitative phenotyping of plant resistance using chlorophyll fluorescence image analysis. Plant Methods, 9(1), 17.

Ruban, A. V. (2016). Nonphotochemical Chlorophyll Fluorescence Quenching: Mechanism and Effectiveness in Protecting Plants from Photodamage. Plant Physiology, 170(4), 1903–1916. doi:10.1104/pp.15.01935

Rungrat, T., Awlia, M., Brown, T., Cheng, R., Sirault, X., Fajkus, J., … Tester, M. (2016). Using phenomic analysis of photosynthetic function for abiotic stress response gene discovery. The Arabidopsis Book, e0185.

Spokas, K., & Forcella, F. (2006). Estimating hourly incoming solar radiation from limited meteorological data. Weed Science, 54(1), 182–189. doi:Doi 10.1614/Ws-05-098r.1

Stowe, B. B., & Yamaki, T. (1957). The history and physiological action of the gibberellins. Annual review of plant physiology, 8(1), 181–216.

Tabas-Madrid, D., Méndez-Vigo, B., Arteaga, N., Marcer, A., Pascual-Montano, A., Weigel, D., … Alonso-Blanco, C. (2018). Genome-wide signatures of flowering adaptation to climate temperature: Regional analyses in a highly diverse native range of Arabidopsis thaliana. Plant, cell & environment.

van Rooijen, R., Aarts, M. G., & Harbinson, J. (2015). Natural genetic variation for acclimation of photosynthetic light use efficiency to growth irradiance in Arabidopsis. Plant Physiology, 167(4), 1412–1429.

Wang, Q. X., Zhao, H., Jiang, J. P., Xu, J. Y., Xie, W. B., Fu, X. K., … Wang, G. W. (2017). Genetic Architecture of Natural Variation in Rice Nonphotochemical Quenching Capacity Revealed by Genome-Wide Association Study. Frontiers in Plant Science, 8. doi:10.3389/fpls.2017.01773

Weigel, D., & Mott, R. (2009). The 1001 genomes project for Arabidopsis thaliana. Genome biology, 10(5), 107.

Wintersinger, J. A., & Wasmuth, J. D. (2015). Kablammo: an interactive, web-based BLAST results visualizer. Bioinformatics, 31(8), 1305–1306. doi:10.1093/bioinformatics/btu808

Zhang, X., Hause, R. J., & Borevitz, J. O. (2012). Natural Genetic Variation for Growth and Development Revealed by High-Throughput Phenotyping in Arabidopsis thaliana. G3-Genes Genomes Genetics, 2(1), 29–34. doi:10.1534/g3.111.001487

