## Supplemental Figures for "A Genome-Wide Association Study of Non-Photochemical Quenching in response to local seasonal climates in *Arabidopsis thaliana*"

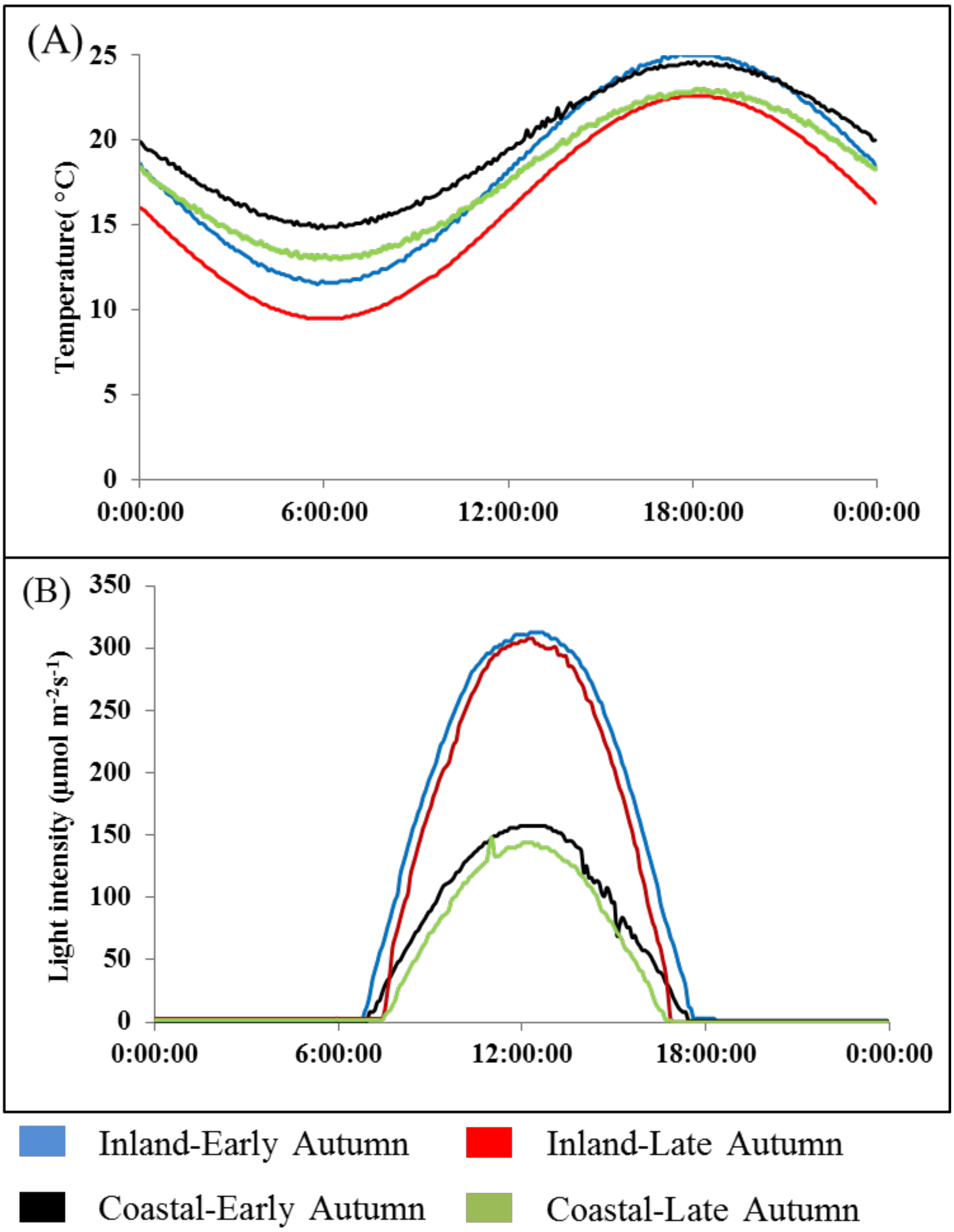


**S1.** Model of simulated Coastal and Inland conditions in early- and late-autumn showing dynamic changes in temperature (A) and light intensity (B

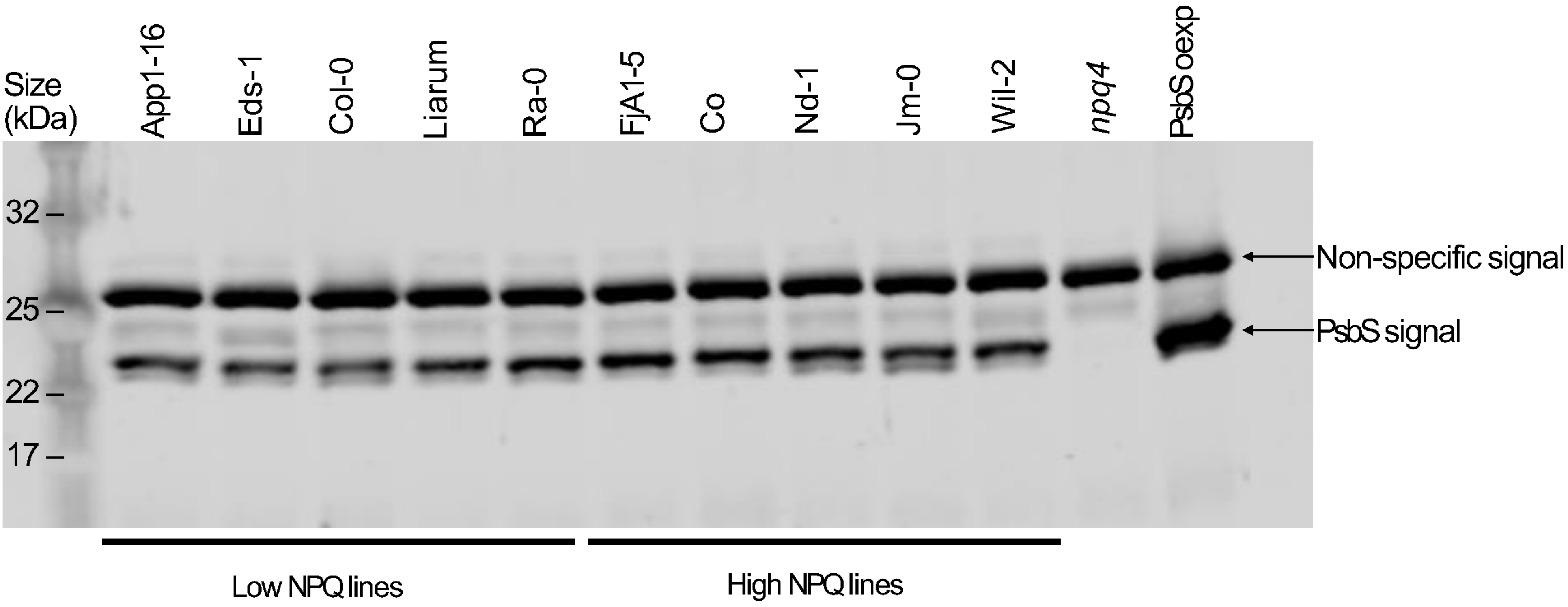


**S2.** Sample western blot showing the PsbS protein content of five low- and high-NPQ plant lines, as well as the *npq4* and PsbS overexpression mutants. The non-specific signal was consistent across all measurements and used to quantify the amount of PsbS protein present in the plant leaf tissue

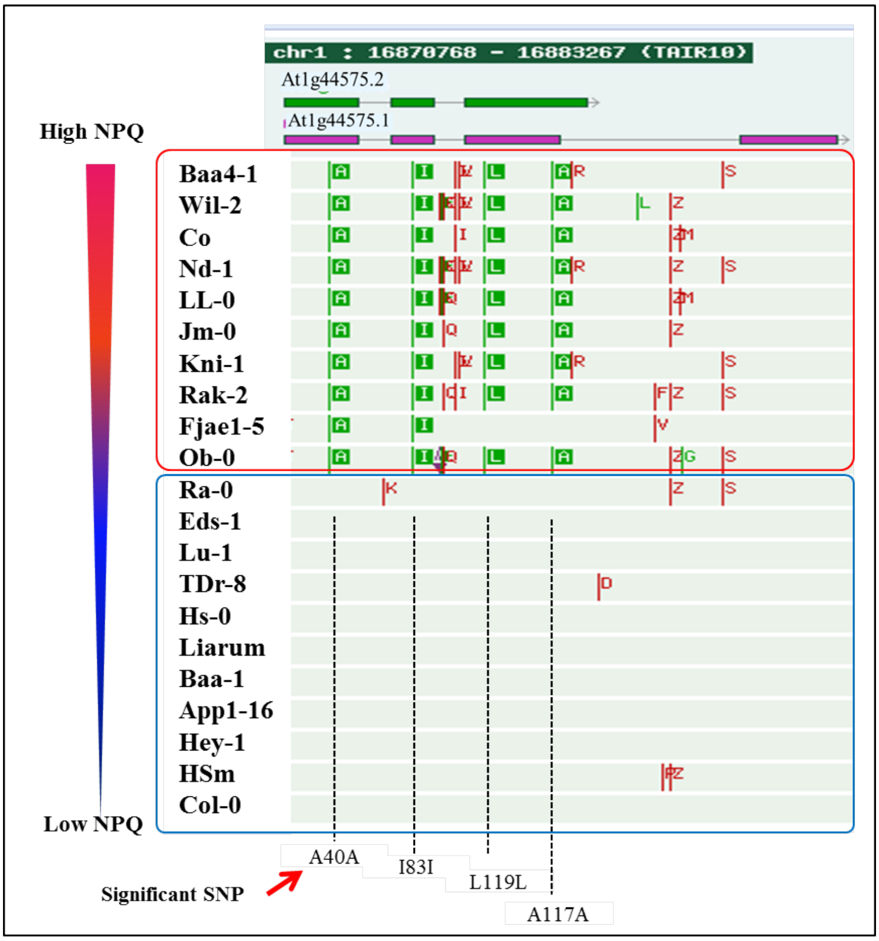


**S4.** Haplotype mapping of high and low NPQ accessions showing amino acid change at the SNPs peaks: synonymous amino acids showed as green colour. Data was generated using online software: *http://signal.salk.edu/atg1001/3.0/gebrowser.php*.

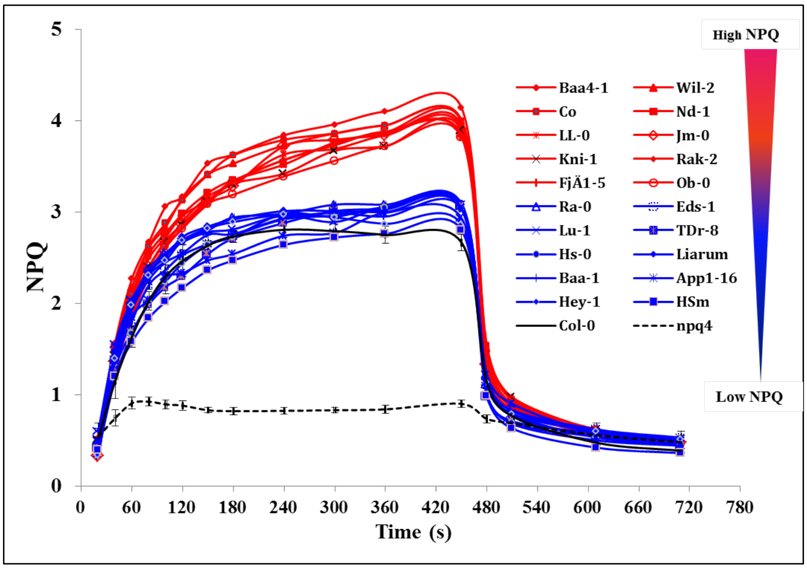


**S6.** NPQ induction of high (red) and low (blue) NPQ haplotypes under Late Autumn Coastal conditions at the 16 leaves stage. The mean + S.D. is given for Col-0 (n=16) and *npq4* (n=4) with the other genotypes represented by one plant.

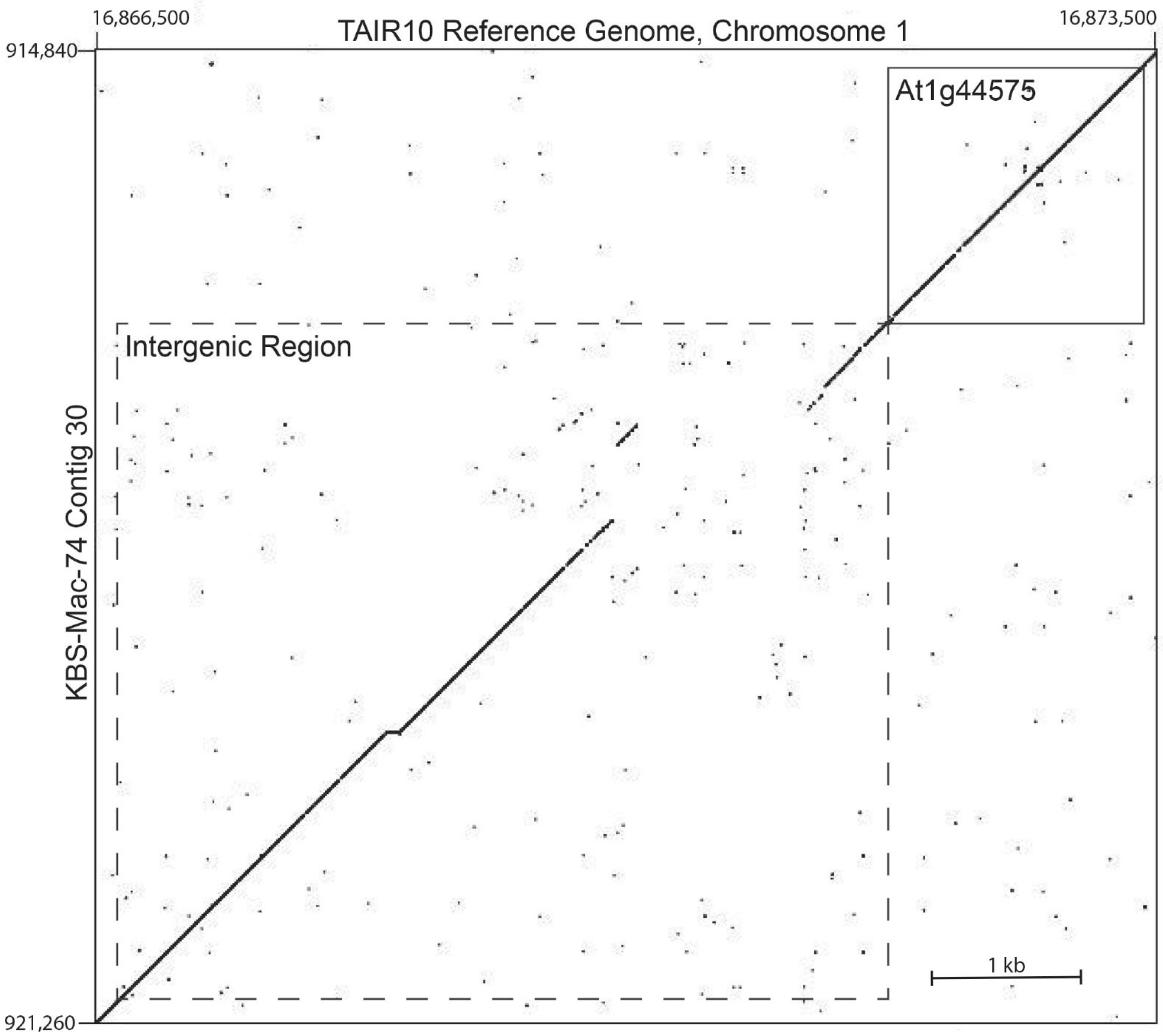


**S7.** Dot matrix comparing *PsbS* genomic regions of TAIR 10 Col-0 Reference Genome and KBS-Mac-74. Col-0 is representative of low NPQ accessions and KBS-Mac-74 is representative of a high NPQ accessions. The *PsbS* gene (At1g44575) and the intergenic regions of the respective genome alignment are labelled with solid-line and dashed-line boxes, respectively. The obvious discontinuity in the intergenic region may indicate the causative agent for the NPQ differences observed in this study.
